# Interphase chromatin as a self-returning random walk: Can DNA fold into liquid trees?

**DOI:** 10.1101/413872

**Authors:** Kai Huang, Vadim Backman, Igal Szleifer

**Affiliations:** Department of Biomedical Engineering, Northwestern University, Evanston, IL 60208, USA.; Chemistry of Life Processes Institute, Northwestern University, Evanston, IL 60208, USA.

## Abstract

We introduce a *self-returning random walk* to describe the structure of interphase chromatin. Based on a simple folding algorithm, our *de novo* model unifies the high contact frequency discovered by genomic techniques, and the high structural heterogeneity revealed by imaging techniques, which two chromatin properties we theoretically prove to be irreconcilable within a fractal polymer framework. Our model provides a holistic view of chromatin folding, in which the topologically associated domains are liquid-tree-like structures, linked and isolated by stretched-out, transcriptionally active DNA to form a secondary structure of chromatin that further folds into a “3D forest” under confinement.

## Introduction

Interphase chromatin is an elephantine biopolymer, which exhibits many exotic properties that are alien to the common sense of traditional polymer physics. The textbook view of interphase chromatin folding, involving 30nm fibers^1^ and their higher-order hierarchical structures, has been greatly challenged by recent ChromEMT experiment^2^, which revealed a highly disordered packing and heterogeneous crowding of DNA. This structural heterogeneity of chromatin has been suggested by Partial Wave Spectroscopy (PWS)^3^, a label-free technique, to have a profound impact on the global transcriptional profile of live cell^4^. To date, different imaging techniques from STORM^5^ to FISH^6^, with various markings, have all demonstrated the heterogeneous and clustering^7^ nature of interphase nuclear materials including DNA, transcription factors and chromatin architecture proteins^8^. On the other hand, it is becoming well known that chromatin exhibits frequent genomic contacts across its 1D contour, largely enhanced inside the topological associating domains (TADs)^9^, first discovered by Hi-C techniques^10^. However, genomic information and imaging data do not easily translate into each other, which poses a fundamental experimental challenge for the comprehensive understanding of chromatin folding. While modeling has been helpful in bridging the gap between the 1D genomic contacts and the 3D chromatin structure^11–21^, most models are highly coarse-grained and focused on local segments of chromatin, given the prohibitive size and complexity of genome. Therefore, it remains a great challenge to model the architecture and crowding effect^22–25^ of the whole macromolecule at a global level without losing the nano-scale details of diverse transcriptional states as shown in Fig. 1A. Lacking a holistic relation between the 1D genomic interactions and the 3D structural heterogeneity, it is still an open question if there is a simple and universal folding principle for chromatin to govern its genomic accessibility and transcriptional activity.

As an alternative to the classical textbook folding picture, it has been hypothesized that chromatin and its encompassing nuclear media are fractal-like^26–30, 22^, with their folding and structure being self-similar across many length scales. In this paper we will theoretically prove that, in the framework of fractal polymer, there is a fundamental yet long-neglected conflict between high spatial heterogeneity and high contact probability. To reconcile this dilemma, we will introduce the concept of self-returning random walk. Within this new framework, we will show that high contact frequency and structural heterogeneity emerge simultaneously. We will demonstrate how SRRW folds a 1D chain into a 3D network of loops and clusters, and discuss the possible molecular mechanisms and biological consequences of our hypothesized chromatin folding.

**Figure 1.**
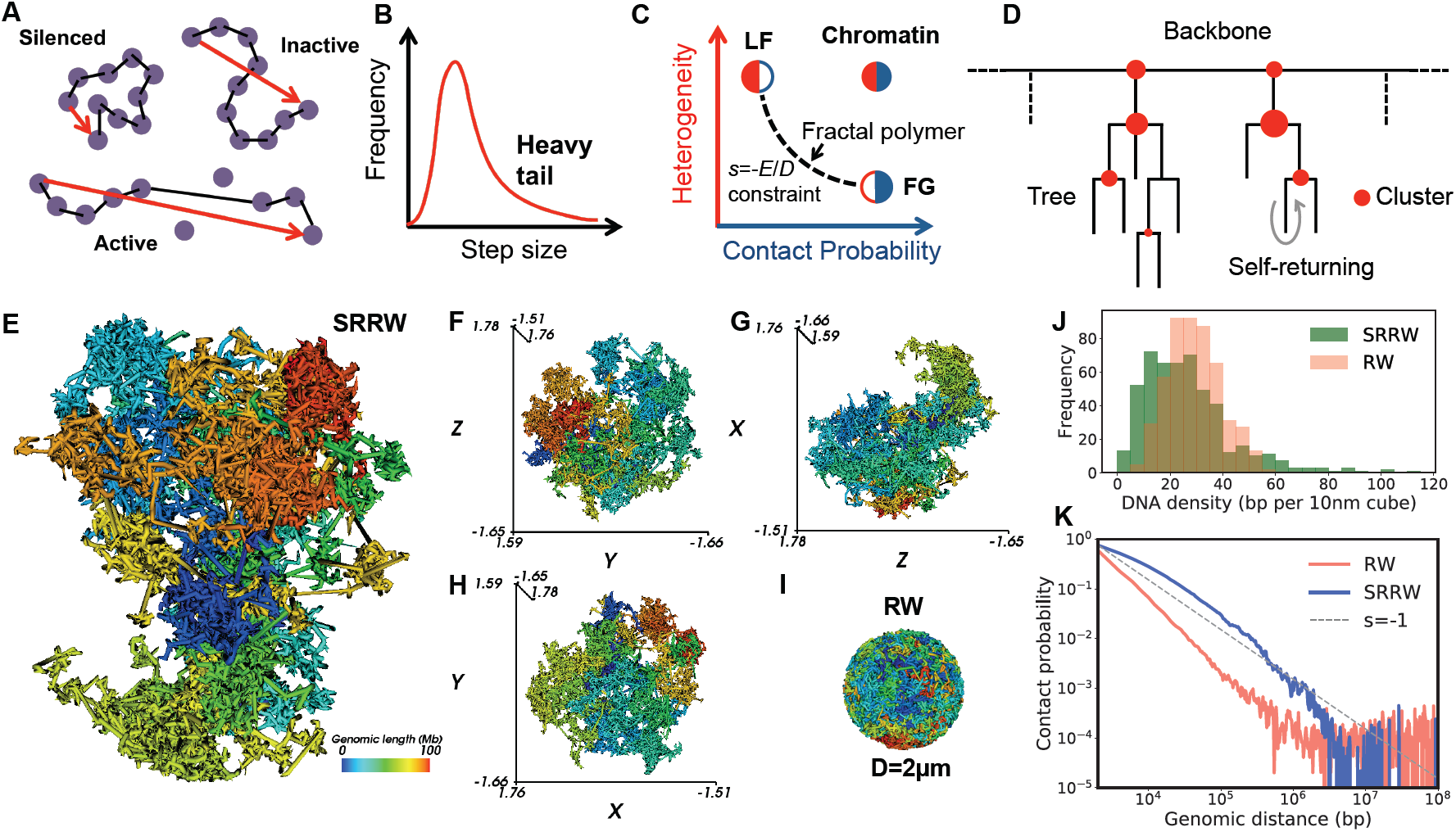
Basic ideas and output of SRRW. (**A**) Diverse epigenetic states at nano-scale. One step approximately maps to 10 nucleosomes or 2kb DNA. (**B**) Modeling different transcription activities by a heavy-tailed distribution of step size. (**C**) Chromatin cannot be a fractal polymer. (**D**) Topology of the architecture of SRRW. (**E-H**) Typical confined SRRW trajectory in 3D and its xyz projections. (**I**) Typical confined RW or EG trajectory. (**J**) Local density spectrums of SRRW and RW sampled by a 150nm-radius probe along the contours. (**K**) Contact probability curves of SRRW and RW.

### Is chromatin a fractal polymer?

Whether chromatin is a fractal^31^ is an intriguing question. Being self-similar allows simple algorithms to generate complicated structures. Given the highly disordered nature of chromatin observed in experiments, it is tempting to describe chromatin conformation in the language of random walk (RW)^32^, whose trajectory is naturally disordered and self-similar. Lévy flight (LF)^33^ as a generalized random walk with a small fractal dimension *D* (*D*<2) seems like a good candidate since it generates a heavy tail of step size (Fig. 1B) that could describe, in a coarse-grained way, a wide range of genomic states, from completely silenced to highly active as illustrated in Fig. 1A. It is also well known that Lévy statistics exhibits clustering at all scales, which could be potentially mapped to the hierarchical sub-TAD/TAD/meta-TAD^34, 35^ structures of chromatin. However, it is not obvious if the highly heterogeneous LF trajectory could facilitate the formation of frequent long-range contacts. On the other hand, the fractal globule (FG)^26, 36^ model with *D*=3 has been widely used to explain the compartmentalization of chromatin and the high contact frequency across all the scales with a power law decay of exponent s=-1. However, FG fills the 3D space in a homogeneous way without forming any clusters. Note that there is a subtle yet important difference between compartmentalization and clustering: clusters are spatially isolated compartments. In other words, compartmentalization is a weaker property than clustering, and without isolation it cannot render density heterogeneity in space. Mounting experimental evidence has revealed that chromatin and its folding agents are rich in clusters^7,8,37^, putting the FG model into question. However, FG is just one extreme case of fractal polymers. Within the infinitely large structural space of fractal polymers, can we find a solution that simultaneously possesses high contact frequency and high structural heterogeneity? Here, based on the argument of scale-invariance and fractal dimensionality, we prove that the answer is negative.

Assume that a generalized RW embedded in an *E*-dimensional space is of fractal dimension *D*. The contact probability *Pc* of distal loci separated by contour length *L* is expected to follow a power law of component *s*, i.e., *Pc*(*L*) ∝ *L^s^*. We assume the criterion of contact formation to be the two loci being closer to each other than a contact distance *c*, or *c*/*ε* in the measure of scale unit. *ε* Changing the contact criterion will rescale the contact probability proportional to (*c*/*ε*) ^*E*^. Also notice that the contour length of a fractal is scale dependent and should be written as *L*(*ε*) = *L*_0_*ε* ^−*D*^, with *L*_0_ being invariant to scaling. Taking into consideration these two facts, we rewrite the contact probability for *L*_0_ under a criterion distance *c* to be *Pc*(*L*_0_, *c*)= *A*(*c*/*ε*)^*E*^ (*L*_0_*ε*^−*D*^)^*S*^ = *Ac*^*E*^*L*_0_^*S*^*ε* ^−(*E*+*DS*)^ where *A* is a constant. Since *Pc* is scale *ε* invariant, *E* +*DS* as to vanish, leading to the relation between *s* and *D* to be

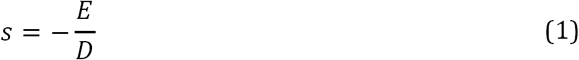

Since *s* and *D* determine the contact frequency and structural heterogeneity of the fractal, respectively, Eq. 1 means that it is impossible for a polymer to simultaneously satisfy (1) high contact frequency, (2) structural heterogeneity and(3) self-similarity. To understand this intuitively: a highly heterogeneous (small *D*) RW fractal such as the LF fills space with self-similar clusters of low inter-cluster contact frequencies, whereas polymer fractals of high contact frequencies, such as the fractal globule, tend to occupy the space in a homogeneous way (large *D*). The fact that interphase chromatin houses both frequent long-range contacts and heterogeneous local DNA densities thus leads us to conclude that its folding structure is not self-similar (Fig. 1C).

### Self-returning random walk

To reconcile the conflict between frequent long-range contacts and high structural heterogeneity, within a RW scheme that respects the disordered nature of chromatin, we introduce a novel type of RW with stochastic, self-returning events. This self-returning random walk (SRRW) organizes its trajectory into clusters of random, tree-like topology, with the inter-cluster contacts boosted by self-returning. The return probability is assumed to decay with the length of the current step size *U*_0_ by a power law:

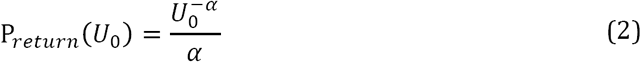

where *α* > 1 is the folding variable for the model chromatin. Smaller *α* leads to higher return frequency. Once a step is returned, a further return is possible based on the same probability function. With a probability of 1-P_return_, a jump from the current position can be issued to explore a new point. The jump takes an isotropic direction and a step size *U*_1_ that follows a power law distribution:

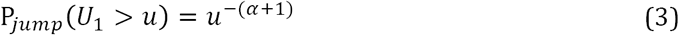

where *u* is larger than 1, the smallest step size in reduced units. We cut off the longest 0.1 percent of the step size so that each jump will not be unrealistically long. To model the confinement effect inside nucleus, we also introduce a global cutoff during the trajectory generation.

An accumulated stack of points track the location of each segment of chromatin, while a subset of them, tracking the unreturned steps, constitute the “backbone” of chromatin. By adding returning to jumping, the model thus turns a chain-generating RW into a network-generator with the degree of networking controlled by the folding parameter *α*. As shown in Fig. 1D, the topological architecture of SRRW is a string of trees with the branches formed by the low-frequency returning of long steps and the nodes formed by the clustering of high-frequency returning of short steps. There is, however, no fine line between small loops and clusters. Isolated by the unreturned long backbone segments, the trees integrate nested loops and clusters into domains for co-regulation.

In our model, we choose each single step to cover 2kb of DNA, which roughly involves an average of 10 nucleosomes. We found such level of coarse-graining appropriate as it renders a good resolution and allows the assumption of non-Gaussian step size that could represent diverse genomic states to hold (Fig. 1A). Based on this genomic length of a single step, we choose the smallest step size to be 30nm long to get a reasonable DNA density of interphase chromatin. We use 50,000 steps to model 100Mb of DNA, roughly the average genomic size of a single chromosome of human. We found an *α* around 1.15 to generate structures that best resemble interphase chromatin, therefore we use *α* = 1.15 in the rest of the paper.

### High contact frequency in the context of high structural heterogeneity

Fig. 1E shows a typical chromatin structure modeled by a confined SRRW, of 100Mb at a 2kb resolution, colored based on the 1D genomic sequence. The long model DNA is clearly folded into domains and compartments reflected by the unmixed color. In contrast, a confined RW, also referred as equilibrium globule, shows no sign of territorial organization (Fig. 1I). On top of the compartmentalization, SRRW is laden with tightly compacted clusters isolated by less compacted DNA segment. As shown in Fig. 1J, SRRW exhibits a wide and non-Gaussian spectrum of local DNA densities or crowding levels, in stark contrast to the case of RW. It is worth noting that the global DNA density heterogeneity of SRRW at micron-scale (Fig. 1E) is a result of the local diversity of transcription activity at nano-scale (Fig. 1A). Such structural coupling across many length scales allows local transcription to shape the global structure of chromatin. The global shape or the territory of SRRW is highly irregular, in sharp contrast with the spherical shape of the FG, but in resemblance with the single-cell experimental findings^38–40^. The shape of SRRW is also highly random, like that of chromosome territory^41^ revealed by single-cell experiment to be cell-to-cell variant^39^. The non-spherical shape of SRRW with large surface area would facilitate high transcription activity inside interchromasomal domain between chromosome territories^42^. The rich porosity of SRRW also allows the accommodation of transcription factories^43^ in the inner space of chromatin^38^.

The distinct folding structures between SRRW and RW give rise to disparate contact probability curves as shown in Fig. 1K. The overall contact frequency of SRRW is much higher than that of RW, except for at the largest genomic scale (above 10Mb). This allows a chromatin folding to avoid unwanted long-range genomic contacts and to enhance local co-regulation of genes. While the RW contact probability decays with a power law exponent s=-1.5 before hitting a plateau due to the confinement, the SRRW has on average a scaling factor around −1, like the FG, but with more scale-dependent ups and downs. Specifically, the SRRW contact probability decay is very slow at the beginning, with s>-1, an abnormal scaling factor from a polymer point of view. Above 100kb, the SRRW contact probability starts to drop faster, and is followed by another slow decay at multi-Mb scale. The range-dependent non-constant scaling behavior of contact probability predicted by SRRW agrees well with experiment^44^ and reiterates the non-fractal nature of chromatin. The two slow probability decays separated by the fast decay have different underlying physics: at small-scale the trees make frequent intra-tree contacts; at large-scale the chromatin backbone folds into a more compacted higher-order 3D structure with enhanced long-range contacts. Between the slow decays the relatively faster decay in the range of a few Mb signifies a topological transition of the architecture of SRRW, from tree-like branched polymer to a non-branching polymeric string of tree domains. The trees generated by SRRW are of stochastic sizes and shapes but rarely contains more than 1Mb DNA. Therefore the contacts at multi-Mb scale are mostly from different trees and happen at low frequency due to the spatial isolation of trees by the stretched backbone segments.

### Tree domains and local hierarchical organization

Having seen the holistic structure of our model chromatin at single cell level in Fig. 1, we now take a closer look of local tree domains. A typical 3D structure of a 3Mb segment of SRRW (1500 steps) is shown in Fig. 2A. The local segment is clearly folded into a bundle of clusters at nano-scale, in line with recent experimental observation^37^. Allowing the conformation in Fig. 2A to relax with preserved topology offers a conformational set (1000 conformations) on which we can obtain a local contact map featuring TAD-like domains as shown in Fig. 2B. Having the 3D structure and the contact map brought side by side by SRRW, we realize that TADs are hierarchical clusters linked and isolated by stretched DNA. In this example, the 3Mb contact map divides itself into two large domains isolated from each other by long steps. Inside these large domains, similar genomic isolations recur at smaller scales, leading to a hierarchical organization of domains. From a loop point of view, this means that large loops often enclose small loops in a hierarchical manner. Such tree-like nested loops avoid entangling and knotting of long DNA in a crowded and disordered environment. On the other hand, condensed chromatin clusters can meet to form nodes of the tree-like structures and get co-repressed. Our prediction is consistent with the fact that TADs are compacted and hierarchical structures with their boundaries being transcriptional active^45^. The average tree size predicted by our model is about 40kb, much smaller than the 200kb average TAD size approximated by early Hi-C experiments^9^, but consistent with recent highresolution TAD analyses^46, 47^. It is worth noting that trees are similar but not equivalent to Hi-C TADs: while Hi-C TADs are loop domains sampled over millions of cells, trees generated by SRRW model are closed loop complexes at a snapshot of time or single-cell level. Recent experiment evidence shows that TADs are dynamic^48^, meaning they are not always closed loops. Given the dynamic nature of loops, we posit that, like chemical reaction, small loops can group into large loops, which can in reverse ungroup into small loops. The contact map in Fig. 2B is based on a frozen topology without any regrouping of loops and therefore could have overestimated the isolation across domains. Nevertheless, as a proof of concept, it clearly demonstrated the fine structure of loop domains and their hierarchical grouping. It has been debated whether higher-order chromatin folding is realized by hierarchical TAD organization^34, 44^. SRRW predicts a hierarchical organization of trees at local scale as we have see in Fig. 2. This hierarchy, however, is not global, as it eventually fades out over scales much larger than the average tree size. If we refer to 10nm fiber as the primary structure of chromatin, SRRW envisions a secondary structure to be a polymeric strand of trees, which further folds into a “3D forest” as the tertiary structure of chromatin. In this folding picture, trees as compounds of nested loops and clusters serve as the functional units to compact DNA and coregulate genes.

**Figure 2.**
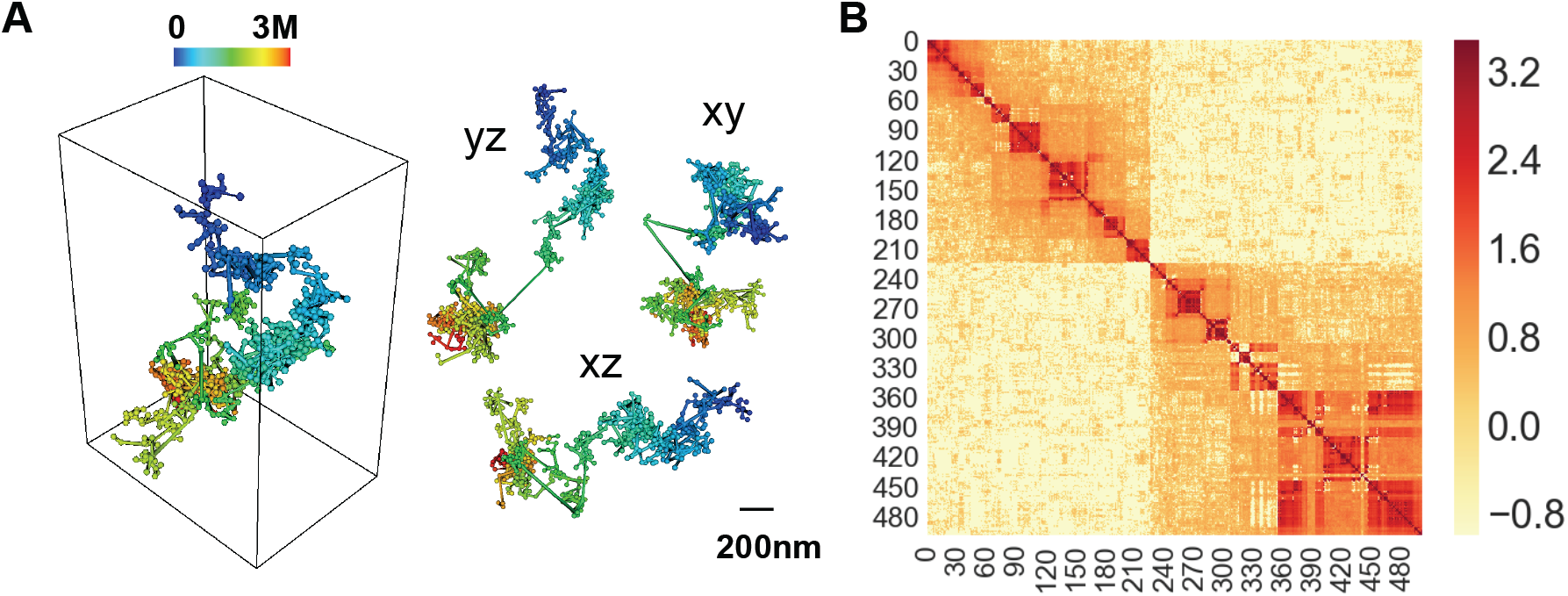
Structure of local SRRW segment and ensemble-averaged contact map. (**A**) 3D structure of a 3Mb segment of SRRW and its xyz projections. (**B**) Contact map of the 3Mb SRRW segment over 1000 random conformations with preserved topology. Color bar in logarithm scale.

### Can DNA fold into liquid trees?

The SRRW model predicts a compaction of chromatin into stochastic tree structures in a liquid state. The idea of chromatin folding into a string of liquid trees might sound counterintuitive, given the 1D topology of DNA and 10nm fiber. However, folding non-branching 10nm fiber into branching higher-order structures could be the key to pivot many potentially conflicting chromatin properties, such as untangled compaction, density heterogeneity, frequent genomic contacts, disordered morphology, and hierarchical domains. Especially, as we have shown that high contact probability and high structural heterogeneity are incompatible for 1D non-branching fractals (Eq. 1), nontrivial topology is needed to solve the conundrum. From a function point of view, tree-like structure is indeed a natural choice to organize large amount of genomic information in a searchable way. However, a fully hierarchical structure would be hard to reprogram, and thus unsuitable for biological evolution in a dynamic environment. Therefore, a balance between local tree structure and global random polymer structure, as predicted by SRRW, could be useful for the genomic “learning”, which optimizes the biological fitness by combining the largely random exploration of the vast genomic landscape (random long-range genomic contacts) and the memorization of fitting genomic contacts (turning long-range inter-tree contacts into local intra-tree contacts during co-replication).

A non-branching polymer without specific guidance will not self-assemble into a tree structure just by thermodynamic processes such as phase separation and gelation. However, DNA can use its sequence code to guide its self-organization and pay an energy price to make it fast. It has been hypothesized that chromatin loops are extruded^49, 50^ by cohesin, a motor protein, in cooperation with CTCF proteins. Since large loops alone have limited compacting capacity, it will be exciting to investigate whether other protein agents could cooperate with CTCF to form nested loops (Fig. 3A). The branches or the self-returned long steps in the tree domains predicted by SRRW could also be realized by transcription-driven supercoiling^51^. To fold into a tree structure, nested or branched supercoiling (Fig. 3B) might be needed. Like loop extrusion, supercoiling requires energy input. But unlike extrusion that requires long-distance travel to form local contacts, supercoiling has the advantage of converting local transcriptional torsion into global structural change in a time-efficient way. Finally, merge of chromatin clusters into tree nodes can also lead to branched structure as shown in Fig. 3C. In fact, clustering and branching are often paired up in SRRW. Given the biological complexity of chromatin, it would not be surprising if multiple folding mechanisms co-exist. The question is how they cooperate and interplay.

**Figure 3.**
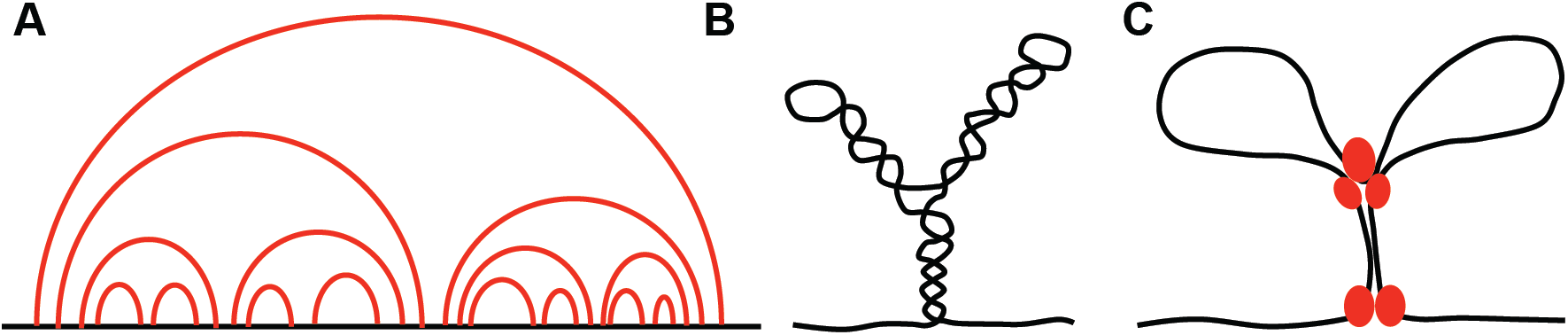
Schematic illustration of some possible folding mechanisms. (**A**) Nested looping in a contour representation (loops are shown in red). (**B**) Branched supercoiling. (**C**) Merge of clusters.

### Conclusion and outlook

We have introduced a new type of RW with self-returning to serve as a minimal physical model of interphase chromatin structure. The SRRW generates conformations that satisfy both high space-filling heterogeneity and high self-interaction frequency, which two properties we have proven to be irreconcilable in the theoretical framework of fractal polymer. It is remarkable that a simple *de novo* model can pivot a wide array of chromatin features, including untangled compaction, disordered morphology, high structural heterogeneity, high contact frequency, and hierarchical genomic domains. This encouraging fact suggests the existence of simple and universal folding principle of interface chromatin. Above 10nm chromatin fiber, SRRW predicts that chromatin is packed into tree-like domains connected by open DNA segments. As the functional units of our model chromatin, trees organize local genomic materials into nested loops and co-repress distal DNA clusters into the tree nodes. Such hierarchical local organization transitions into a worm-like secondary structure, which is highly flexible for further global 3D packing without much concern of entangling and knotting due to its reduced contour dimension. Under confinement, the secondary structure folds into a “3D forest” with enhanced long-range contacts, large surface area and great porosity. Our model highlights non-Gaussian statistics of chromatin structure and emphasizes the fact that chromatin is a living polymer whose folding is likely to be out-of-equilibrium and energy consuming. As a new type of RW, we hope SRRW will find useful applications in a variety of fields concerning living systems and networks.

## Acknowledgements

We gratefully acknowledge funding from National Science Foundation Grants: Biol & Envir Inter of Nano Mat 1833214, EFRI RESEARCH PROJECTS 1830961.

